# Gap junctions in the alimentary tract regulate reproductive span in *C. elegans*

**DOI:** 10.64898/2026.06.03.729750

**Authors:** Hitomi Hidaka, Kazuma Akashi, Shiying Wang, Shohei Yamamoto, Takumi Chinen, Shoji Hata, Masamitsu Fukuyama, Daiju Kitagawa

## Abstract

Aging does not occur uniformly throughout an organism but is instead differentially regulated across distinct physiological systems. In particular, reproductive aging is often temporally distinct from somatic aging and exhibits species-specific trajectories, suggesting that different physiological functions may age independently. Here we show that mutation of *inx-20*, an innexin family gene encoding a gap junction component, markedly extends reproductive span, with only a minor increase in overall lifespan. Furthermore, this extension of reproductive span persists in a feminized genetic background, thereby precluding the possibility that it is driven by altered sperm dynamics. *inx-20* is expressed in a specific subset of cells within the alimentary tract, and its expression is selectively repressed in a fraction of these cells during dauer diapause, suggesting a role in nutrient responses. Genetic analyses suggest that *inx-20* operates via a distinct mechanism that does not intersect with the TGF-β and IIS-FOXO pathways, which are established regulators of reproductive span. Collectively, our results suggest that gap junctions in the alimentary tract are a selective determinant of reproductive span, capable of extending it substantially without a commensurate effect on lifespan.

## Introduction

Aging does not proceed uniformly across the whole body, but rather occurs at different rates across each physiological function. In particular, reproductive functions age independently of other bodily functions and exhibit species-specific patterns of aging (Jones *et al*., 2014). In human females, fertility declines earlier and over a more compressed time window than many physiological functions: while motor and cognitive functions decline gradually across decades of adult life, reproductive function declines over roughly a decade from the mid-30s and is almost completely lost before menopause (Te Velde and Pearson, 2002; Salthouse, 2009; Furrer and Handschin, 2025).

Similar to human females’ reproductive senescence, *Caenorhabditis elegans* exhibits early decline in reproductive capability (Hughes *et al*., 2007). More specifically, self-fertilizing worms cease reproduction within approximately one week and survive for an additional one to two weeks (Luo *et al*., 2009). Suppression of the insulin/IGF-1 signaling (IIS) pathway and TGF-β signaling pathway, as well as dietary restriction induced by *eat-2* mutation have been shown to extend reproductive span: the time from the onset of egg laying to the end of progeny production (Hughes *et al*., 2007; Luo *et al*., 2009, 2010). These pathways were identified through previous studies on lifespan regulation. In addition, relatively few systematic genetic screens have been conducted to identify genes that regulate reproductive span. One of these screens, a forward genetic screen with clonal F_1_ isolation, identified the *phm-2* mutant, which exhibits extension of both lifespan and reproductive span (Hughes, Huang and Kornfeld, 2011). This mutant exhibits bacterial colonization of the intestine and subsequent activation of innate immunity, which induces bacterial avoidance and thereby promotes dietary restriction-like effects that contribute to reproductive span extension (Kumar *et al*., 2019). Reproductive span measurement requires complex handling, including frequent transfer of worms, making it difficult to perform large-scale classical clonal F_1_ screens for mutants with extended reproductive span (Jorgensen and Mango, 2002). Indeed, the screen conducted by Hughes et al. was limited to approximately 1,000 F_1_ animals (Hughes, Huang and Kornfeld, 2011). The other conducted a genome-wide RNAi screen to identify approximately 30 genes which are related to IIS–FOXO pathway or TGF-β Sma/Mab pathway (Wang *et al*., 2014). RNAi efficiency in *C. elegans* can vary depending on tissue, developmental stage, target gene, and genetic background (Simmer *et al*., 2002; Wang *et al*., 2005; Lehner *et al*., 2006; Schmitz, Kinge and Hutter, 2007). The screen conducted by Wang et al. used an RNAi-hypersensitive strain to enhance RNAi potency (Wang *et al*., 2014). While RNAi-hypersensitive strains are useful for increasing RNAi penetrance, their underlying mutations are not phenotypically neutral (Ceron *et al*., 2007). Thus, despite these important studies, unbiased forward genetic screens for reproductive span regulators remain limited.

Gap junctions, which are composed of connexins in vertebrates and innexins in invertebrates, transfer small molecules such as ions, ATP, and inositol 1,4,5-trisphosphate (IP_3_), and mediate intercellular communication (Evans and Martin, 2002). Through this direct cytoplasmic coupling, gap junctions coordinate tissue-level activities, including electrical signaling, metabolic coupling, and synchronized physiological responses across multicellular tissues (Dbouk *et al*., 2009; Goodenough and Paul, 2009; Simonsen, Moerman and Naus, 2014). In *C. elegans*, innexins have been implicated in neuronal electrical coupling, body-wall muscle coupling, propagation of intercellular Ca^2+^ waves in the intestine, and soma-germline communication required for gametogenesis (Peters *et al*., 2007; Liu *et al*., 2013; Starich, Hall and Greenstein, 2014; Jin *et al*., 2020). A notable feature of connexins and innexins is that their expression dynamically changes in response to environmental conditions, thereby modulating intercellular connectivity through gap junctions. For example, in the vertebrate retina, connexin 36-mediated coupling is modulated by circadian rhythm and the light environment to dynamically regulate visual sensitivity and network state (Zhang *et al*., 2020). In *C. elegans*, expression of *inx-6* is induced in the AIB interneuron during dauer and L1 diapauses, leading to remodeling of locomotory behavior (Bhattacharya *et al*., 2019). Connexin and innexin expression is also altered in tissues other than neurons, but their physiological functions remain uncharacterized.

Here, we developed a large-scale screening method identifying genes that affect reproductive span in which mutagenized worms are handled in bulk rather than individually handpicked. Using this screen, we isolated a loss-of-function mutant of the gap junction component innexin, *inx-20*, which exhibits an extended reproductive span. This mutant shows only a moderate extension of lifespan. *inx-20* is expressed in restricted regions of the alimentary tract, and its posterior expression is specifically lost during dauer diapause. Furthermore, this mutant extends reproductive span through a pathway that is at least partially distinct from the IIS and TGF-β Sma/Mab pathways.

Our findings demonstrate that gap junctions in the alimentary tract, but not in neurons, regulate reproductive span through a pathway that is not fully explained by canonical reproductive longevity pathways.

## Results

### Loss of *inx-20* function extends reproductive span specifically

To identify genes that influence reproductive aging, we established a larger-scale screening strategy to identify genes contributing to reproductive span extension. We separated hundreds of F_2_ adult descendants of mutagenized worms from F_3_ larvae and embryos by sedimentation. The adults were then transferred in bulk onto 10 cm seeded plates every day until day 10 or 11 of adulthood, when most wild-type worms have ceased reproduction. This procedure enabled high-throughput screening of approximately 144,000 haploid genomes (Fig. 1a). From this screen, we found six strains that exhibited extended reproductive span, two of which also showed an extension of mated reproductive span (Fig. S1a). We isolated one strain which exhibits increased late reproduction after outcrossing two times (Fig. S1b). Whole-genome sequencing revealed mutations in four protein-coding genes, including amino acid substitutions and splice-site mutations (Fig. S1c). Of these four genes, *F46A8*.*4* is expressed specifically in males, whereas the other three are expressed in hermaphrodites. To identify the causative mutation, we recreated each of the three mutations found in the isolated strain using the CRISPR-Cas9 system. On adult day five, when more than half of wild-type worms had ceased reproduction, only the mutation in *inx-20* phenocopied the isolated strain with respect to the proportion of reproducing individuals (Fig. S1c). In the following experiments, we confirmed that *inx-20* is the gene responsible for the extended reproductive span.

**Figure 1.**
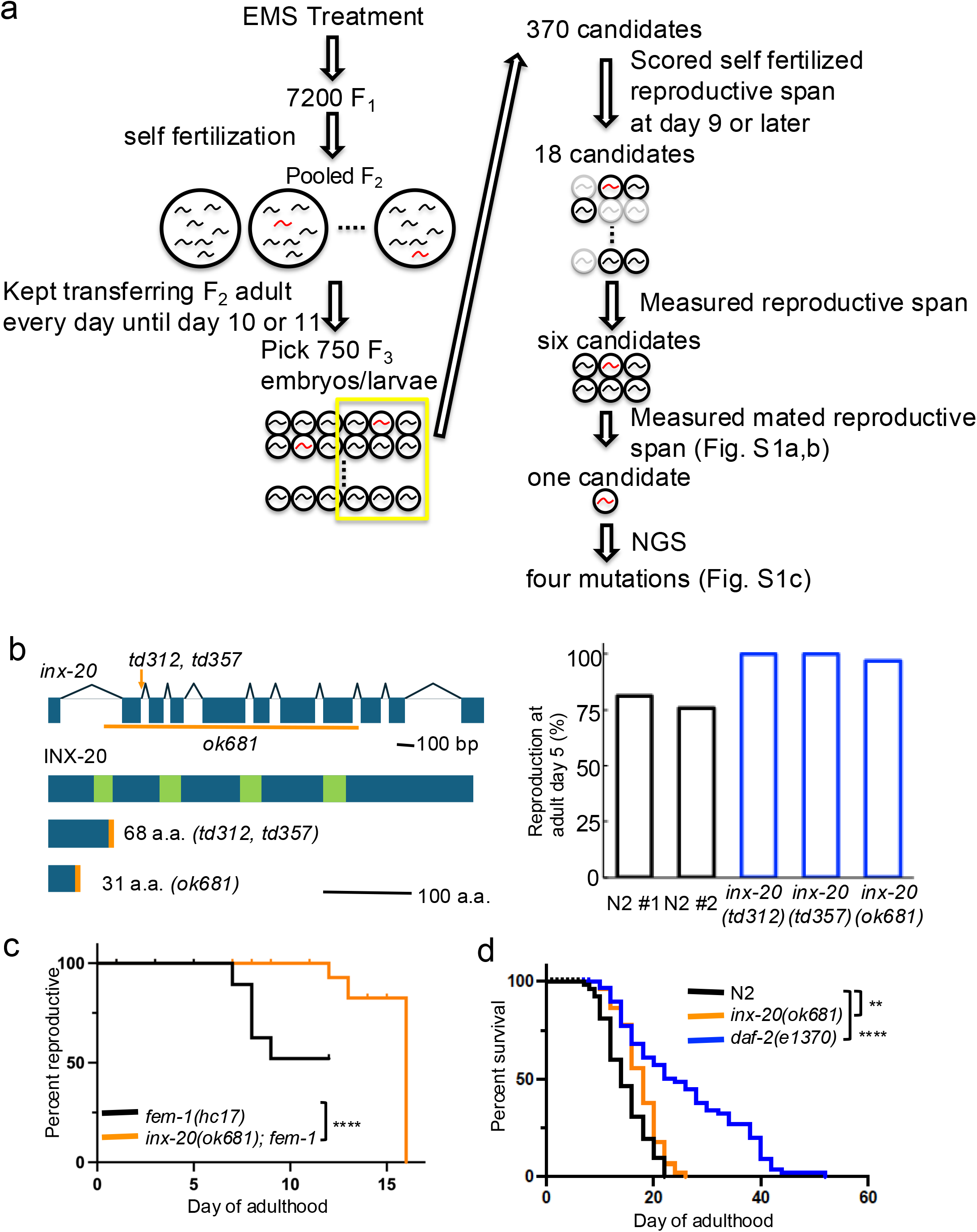
Loss of *inx-20* function extends reproductive span without lifespan extension (a) Schematic of the forward genetic screening strategy used to identify mutants with altered reproductive span. Hermaphrodites derived from mutagenized worms were screened in bulk, and candidates were selected based on extended reproductive span. See Materials and Methods for details. See figure S1a and S1b for mated reproductive span measurement, and figure S1c for NGS-based identification of causal mutations. (b) Schematic representation of the *inx-20* gene and mutant alleles used in this study. Exons are shown as boxes and introns as lines. *td312* and *td357* are nonsense mutations predicted to introduce premature stop codons. *ok681* is a deletion allele. Predicted transmembrane domains are shown in green. *inx-20* point mutation *inx-20(td357)* worms and deletion mutation *inx-20(ok681)* worms phenocopy isolated *inx-20* mutation *inx-20(td312)* worms. The *inx-20(td312)* mutant was examined in experiment #1, whereas the other mutants were analyzed in experiment #2. (c) Reproductive span extension in *inx-20* mutants occurs independently of sperm quality. To suppress sperm production, hermaphrodite worms were maintained at 25 °C until L4 larval stage. (d) *inx-20(ok681)* mutant worms exhibited only a modest extension of lifespan.

The *inx-20* gene consists of nine exons. The *td312* allele is a single-base substitution that introduces a premature stop codon and is predicted to produce a truncated protein (Fig. 1b). To determine whether *inx-20* is responsible for the extended reproductive span, we generated the *td357* allele carrying the same nucleotide change as *td312* and confirmed that both the *td357* and the *ok681* deletion allele recapitulate the *td312* phenotype in terms of reproductive output on adult day five (Fig. 1b). In the following experiments, we used the *ok681* allele as an *inx-20* mutant because of its ease of handling. *inx-20(ok681)* mutant worms reproducibly exhibited a markedly extended reproductive span (Fig. S1d).

Self-fertilized hermaphrodites cease reproduction due to the depletion of self-sperm (Ward and CARREL Carnegie, 1979). To evaluate reproductive span independently of sperm depletion, we generated *inx-20(ok681); fem-1(hc17)* double mutants. *fem-1(hc17)* mutants become spermless when maintained at the restrictive temperature (25 °C) until the L4-stage larvae (Nelson, Lew,’and and Ward’, 1978). Mating *fem-1(hc17)* background mutants with N2 males allowed us to measure reproductive span under sufficient and equivalent sperm conditions. As a result, the extended reproductive span in *inx-20* mutant worms was observed independently of self-sperm depletion (Fig. 1c). This result indicates that loss of *inx-20* in the somatic cells or oocytes, rather than in the sperm, underlies the extended reproductive span. We next tested whether expression of wild-type *inx-20* from an extrachromosomal array could rescue the *inx-20(ok681)* phenotype. A construct expressing C-terminally mVenus-tagged *inx-20* was injected into *inx-20(ok681)* mutants, and transgenic animals were identified by body-wall muscle mCherry expression as a co-injection marker. Using this strain, we successfully rescued the extended reproductive span phenotype in *inx-20* mutant (Fig. S1e). This result supports that the loss of *inx-20* function extends reproductive span. Furthermore, *inx-20* mutants exhibit only a moderate extension in lifespan (Fig. 1d). This phenotype contrasts with that of *daf-2* mutants, which exhibit substantial extension of both reproductive span and lifespan (Luo *et al*., 2009). Together, these results indicate that *inx-20(ok681)* exhibits an extended reproductive span without markedly extending lifespan.

### Dauer-dependent partial loss of *inx-20* expression in the alimentary tract

To understand how *inx-20* regulates reproductive span, it is essential to precisely characterize its spatiotemporal expression pattern. *inx-20* encodes an innexin, a component of gap junctions. To elucidate the expression pattern of *inx-20*, we knocked in mStayGold::AID* tag immediately before the stop codon of the endogenous *inx-20* locus using CRISPR-Cas9-mediated genome editing (see Materials and Methods). INX-20::mStayGold expression was observed in the alimentary tract largely consistent with previous analysis using 1-kb upstream region of the *inx-20* ORF as the promoter sequence (Fig. 2a) (Altun *et al*., 2009). INX-20::mStayGold expression was not detected in comma-stage embryos but became apparent during late embryogenesis (Fig. S2a), and remained detectable through at least adult day five (Fig. S2b). The expression of several innexins in neurons is altered during the dauer stage to facilitate behavioral adaptation. For example, *inx-6* is induced under starved condition such as dauer and L1 diapauses in AIB neurons, and its expression during dauer diapause remodels locomotory behavior. However, *inx-20* expression at the intestinal-rectal valve specifically becomes undetectable during the dauer stage but remains detectable under starvation alone (Fig. 2b). This observation suggests that *inx-20* may function as part of a physiological adaptation program that is actively regulated during dauer diapause.

**Figure 2.**
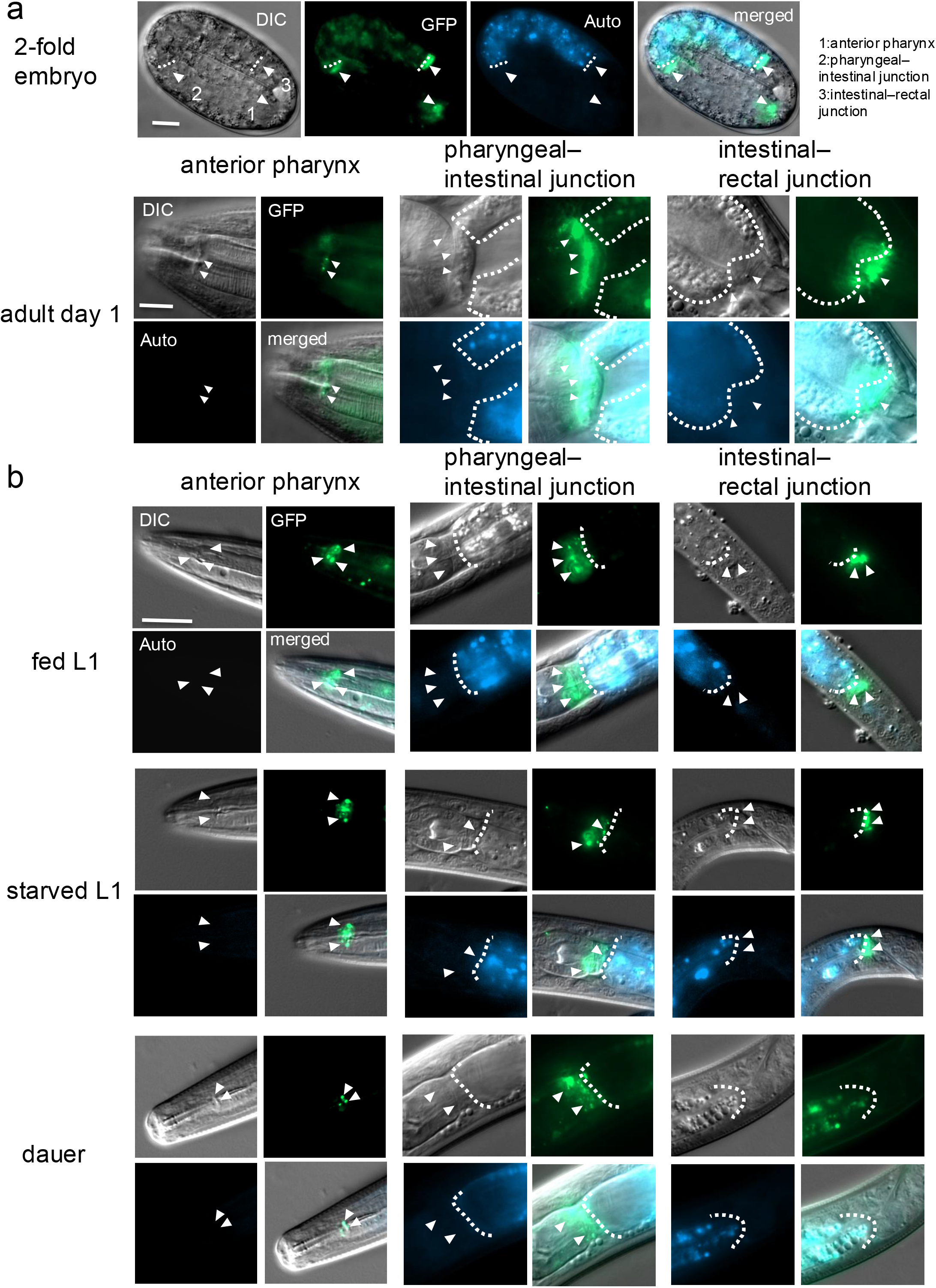
Dauer-dependent partial loss of *inx-20* expression in the alimentary tract (a) Under the standard condition, INX-20::mStayGold(mSG) is continuously expressed from the embryonic stage to adulthood in the anterior pharynx, at the pharyngeal– intestinal junction, and at the intestinal-rectal junction. (b) Expression at the intestinal-rectal junction was no longer detectable specifically in dauer larvae. (a) and (b) INX-20::mSG expression is shown in green (panels labeled GFP), and intestinal autofluorescence is shown in blue (panels labeled Auto). Arrowheads indicate INX-20::mSG expression, and the dotted line marks the intestinal boundary. In all panels except the two-fold embryo, anterior is to the left and posterior to the right. At least 20 worms were examined for each condition, and representative images are shown. Scale bar, 10 µm. Worms were bleached and arrested at the L1-stage. The L4-stage was reached 43.5 h after release, and day one adults were obtained 68.5 h after release. Dauer larvae were prepared by keeping culture plates at 25 °C for several days.

### *inx-20* regulates reproductive span in a pathway independent of the insulin/IGF-1(IIS) pathway and TGF-β Sma/Mab pathway

To determine whether *inx-20* regulates reproductive span through established longevity pathways, we examined its genetic interactions with key reproductive aging regulators. We crossed *inx-20(ok681)* mutant with *daf-2(e1370)* mutant, which encodes the insulin/IGF-1 receptor, and *sma-2(e502)* mutant, which encodes a transcription factor of the TGF-β Sma/Mab pathway, to assess whether the double mutants exhibit a further extension of reproductive span compared with either single mutant.

The *inx-20; daf-2* double mutant did not exhibit a further extension of reproductive span compared to either single mutant (Fig. 3a). The reproductive span extension observed in *daf-2* mutants is dependent on *daf-16/FOXO*, which is a major downstream effector of the IIS pathway, and is completely abolished by loss of *daf-16* (Hughes *et al*., 2007). We next crossed *inx-20(ok681)* mutant with *daf-16(mu86)* mutant to examine the genetic requirement of *daf-16* for the extended reproductive span of *inx-20* mutants. However, the extended reproductive span of *inx-20* mutants was not suppressed by loss of *daf-16*, in contrast to the *daf-16*-dependent reproductive span extension observed in *daf-2* mutants (Fig. 3b). Furthermore, *daf-2(e1370)* mutant larvae become dauer when postembyonically cultured at 25 °C (Riddle, Swanson and Albert, 1981). In contrast, *inx-20(ok681)* mutants neither exhibit this *daf-2* phenotype nor suppress the dauer formation in *daf-2(e1370)* mutants (Fig. 3c). Collectively, our genetic analyses suggest that *inx-20* is unlikely to act within the insulin/IGF-1 signaling pathway that regulates dauer formation and reproductive aging, and that its role in reproductive span regulation is not mediated by *daf-16/FOXO* activity.

**Figure 3.**
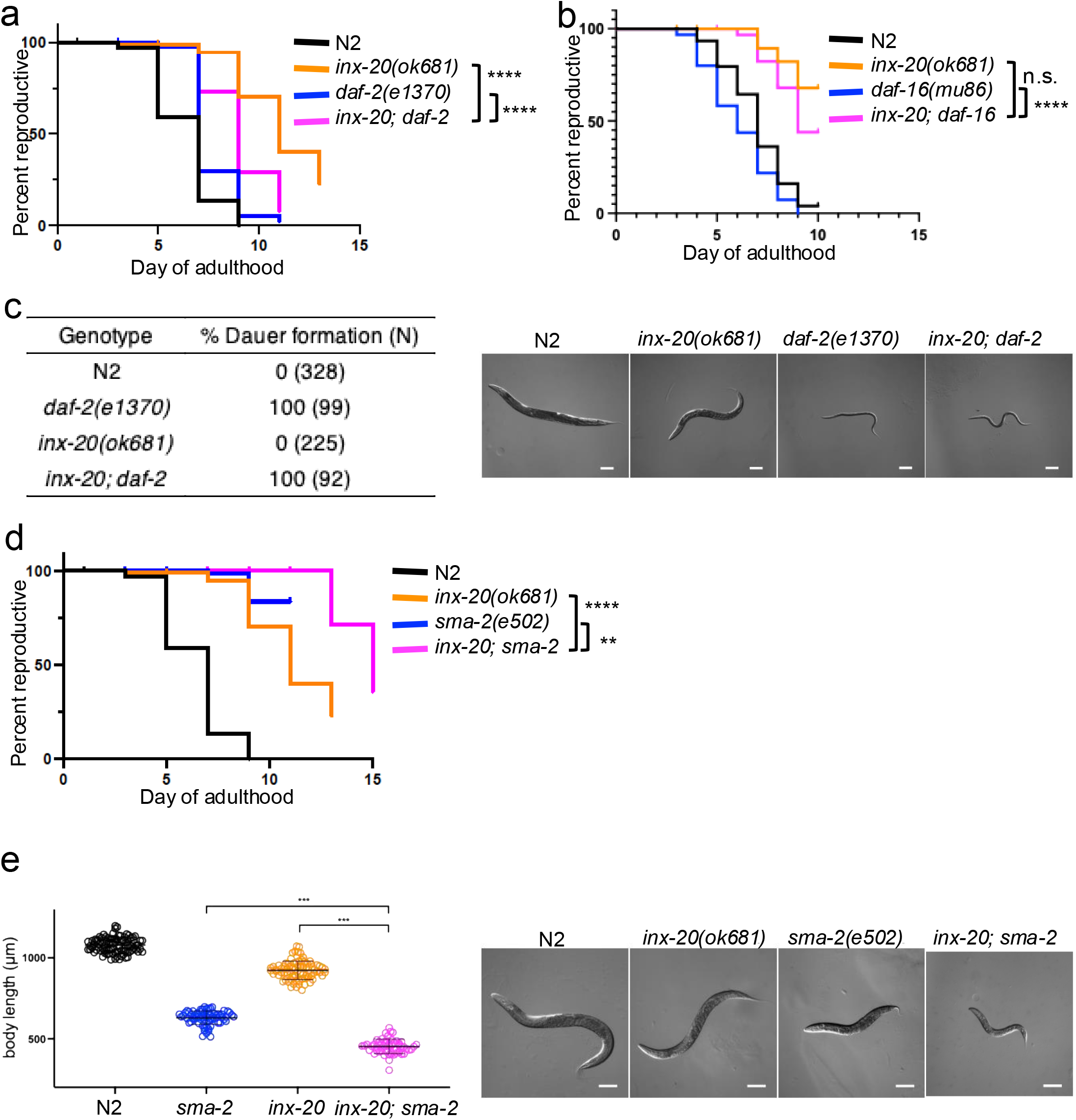
*inx-20* regulates reproductive span in a pathway independent of the IIS pathway or TGF-β (Sma/Mab) pathway (a) The *inx-20(ok681); daf-2(e1370)* double mutants do not show an additive extension of reproductive span relative to either single mutant. (b) The *inx-20(ok681); daf-16(mu86)* double mutants exhibit a reproductive span extension comparable to that of *inx-20(ok681)* single mutants. (c) The *inx-20(ok681)* mutation does not induce dauer entry, nor does it prevent dauer formation in the *daf-2(e1370)* background. (d) The *inx-20(ok681); sma-2(e502)* double mutants exhibit a significant extension of reproductive span relative to either single mutant. (e) *inx-20* and *sma-2* act independently to reduce body size. Scale bar, 100 µm. (a),(b), and (d) Log-rank (Mantel-Cox) test, 30-50 worms per strain, p-value * < 0.05, ** < 0.01, *** < 0.001, **** < 0.0001. (a) and (d) n = 3, (b) n = 1. (e) Mann–Whitney U test (Wilcoxon rank-sum test).

Similar to the *inx-20* mutant (Fig. 1d), loss of *sma-2*/Smad leads to a marked extension of reproductive span, while lifespan extension is modest (Luo *et al*., 2009). However, the *inx-20; sma-2* double mutant displayed a significantly greater extension of reproductive span than either single mutant (Fig. 3d). Furthermore, the *inx-20;sma-2* double mutants are smaller than *sma-2* single mutants (Fig. 3e). Both results suggest that *inx-20* functions in a pathway distinct from TGF-β Sma/Mab signaling.

## Discussion

Here, we report the results of a forward genetic screen for *C. elegans* mutants exhibiting an extended reproductive span. The screen identified *inx-20* as a novel regulator of reproductive aging, as its loss extends reproductive span via a pathway that is independent of previously established regulatory mechanisms (Luo *et al*., 2009, 2010; Hughes, Huang and Kornfeld, 2011; Kumar *et al*., 2019).

Despite the large screening scale (144,000 haploid genomes), our forward genetic screen failed to recover known mutants with extended reproductive spans, including mutants in the IIS and TGF-β pathways, as well as *eat-2* and *phm-2* mutants (Luo *et al*., 2009, 2010; Hughes, Huang and Kornfeld, 2011; Kumar *et al*., 2019), though it successfully identified *inx-20* as a novel factor whose loss extends reproductive span. This is likely because our screening protocol, particularly the step separating adults from F_2_ larvae by sedimentation, biased against mutants with slow growth or small body size, traits often associated with those pathways.

The expression of gap junction proteins, including connexins in vertebrates and innexins in invertebrates, varies considerably across cell types and developmental stages. One of the best-characterized examples is the light-dependent regulation of gap junctions between retinal horizontal cells in vertebrates, where junctional coupling is modulated in response to changes in ambient illumination, thereby adjusting receptive field size and balancing spatial resolution with sensitivity (Bloomfield and Völgyi, 2009). Consistent with the idea that gap junctions can be remodeled in response to environmental cues, we found that expression of *inx-20* at the intestinal-rectal valve is downregulated specifically during the dauer stage. To our knowledge, this is the first demonstration that innexin expression in the *C. elegans* alimentary tract is subject to environmental regulation, extending the concept of dauer-induced gap junction remodeling beyond the nervous system (Bhattacharya *et al*., 2019). Although whether other innexins in the alimentary tract are similarly regulated upon dauer entry remains to be determined, this observation raises the possibility that the *C. elegans* alimentary tract alters its function and activity by remodeling the abundance and composition of gap junctions during diapause. This dauer-associated reduction of *inx-20* expression may also provide a framework for interpreting the extended reproductive span of *inx-20* mutants. Because dauer entry involves broad physiological remodeling and can leave lasting effects on adult gene expression and life-history traits (Hall *et al*., 2010; Ow *et al*., 2018), loss of INX-20 function may partially mimic a dauer-like state in the alimentary tract. Such a state could alter alimentary tract activity or intestine-germline signaling, thereby selectively delaying reproductive aging.

In *C. elegans*, previous studies have demonstrated that the expression of innexin genes in specific neuronal populations is remodeled during the dauer stage, contributing to dauer-specific behavioral changes (Bhattacharya *et al*., 2019). Gap junctions composed of innexins in the alimentary tract enable electrical and Ca^2+^ coupling between cells, thereby contributing to synchronized rhythmic activity and coordinated contractile movements. For instance, *inx-6* regulates pharyngeal pumping, and *inx-6* mutants exhibit feeding defects (Li, Dent and Roy, 2003). In addition, *inx-16* mediates the propagation of Ca^2+^ waves in the intestine, thereby regulating synchronized intestinal oscillations to drive the defecation cycle (Teramoto and Iwasaki, 2006; Peters *et al*., 2007). Although the precise role of the intestinal-rectal valve cells in Ca^2+^ wave propagation and the defecation cycle remains to be determined, downregulation of *inx-20* during dauer diapause may contribute to the suppression of the defecation cycle. This would be consistent with the non-feeding state of dauer larvae, in which food intake is prevented by the sealing of the buccal cavity by a cuticular block, and pharynx pumping is completely suppressed (Cassada and Russell, 1975; Popham A N and Webster, 1979). If similar remodeling is mimicked in *inx-20* mutants, altered alimentary tract motility could change nutrient flux or availability and thereby induce a dietary restriction-like state. Because dietary restriction, including the *eat-2* model of reduced feeding, delays reproductive aging and extends reproductive span in *C. elegans* (Lakowski and Hekimi, 1998; Hughes *et al*., 2007), such metabolic changes may contribute to the extended reproductive span of inx-20 mutants.

*inx-20* expression is restricted to discrete regions at both ends of the intestine (Fig. 2a), and mutants exhibit a marked extension of reproductive span despite showing only modest changes in lifespan (Fig. 1c,d). These observations suggest that *inx-20* may primarily affect reproductive aging rather than somatic aging, potentially weakening the coordination between these two aging processes. Aging can be viewed as a process in which functional decline occurs progressively and coordinately across multiple tissues. In *C. elegans*, correlations have been reported between lifespan and age-associated declines in locomotor and feeding activities (Huang, Xiong and Kornfeld, 2004). Consistent with this view, studies in *C. elegans* have shown that aging-related phenotypes can be shaped by both tissue-specific and inter-tissue mechanisms (Miller *et al*., 2020). Thus, *inx-20* may contribute to a mechanism that links alimentary tract function with reproductive aging, although the relevant cellular interactions remain to be determined. One possible anatomical context for this link is the posterior enteric region, where defecation and egg-laying circuits are closely associated. Intestinal Ca^2+^ oscillations are initiated from the posterior intestine (Espelt *et al*., 2005), and *inx-20* is expressed near the sphincter muscle, part of the enteric musculature activated by AVL/DVB motor neurons during the defecation expulsion step (Mclntlre *et al*., 1993). AVL/DVB neurons are functionally connected with the HSN egg-laying circuit, which induces the egg-laying active phase (Waggoner *et al*., 1998), and defecation and egg-laying circuits are coordinately regulated through chemical and electrical synapses (Garcia and Collins, 2022). Based on this anatomical and functional relationship, altered *inx-20* function could influence egg-laying rate and reproductive span indirectly through the defecation-associated enteric circuit.

## Acknowledgements

We thank all members of Kitagawa laboratory for fruitful discussion.

This work was supported by Japan Society for the Promotion of Science (JSPS) KAKENHI grants 18K06246, 22H02629, 22K19305 (MF), 22K20624, 23K14176 (SY), 20K15987, 23H02627 (TC), 20K22701, 21H02623, 22K19370, 24K02174 (SH), 19H05651, 24H02284 (DK), Japan Science and Technology Agency (JST), the PRESTO program JPMJPR21EC (SH) Japan Science and Technology Agency (JST), the CREST program JPMJCR22E1 (DK), AMED-PRIME 21gm6110017h0004 (MF), Takeda Science Foundation (DK, SH, TC, SY), Uehara Memorial Foundation (SH, TC, SY), Research Foundation for Pharmaceutical Sciences (SH, TC, SY), Koyanagi Zaidan (MF), Mishima Kaiun Memorial Foundation (MF), Kanae Foundation for the Promotion of Medical Science (TC), Kato Memorial Bioscience Foundation (SH), Naito Foundation (TC), Heiwa Nakajima Foundation (TC), Sumitomo Foundation (TC), Inamori foundation (SY) Astellas Foundation for Research on Metabolic Disorders (SY).

Some strains were provided by the CGC, which is funded by NIH Office of Research Infrastructure Programs (P40 OD010440)

## Author contributions

Conceptualization: H.H., M.F. and D.K.; Methodology: H.H. and M.F.; Formal analysis: H.H. and K.A.; Investigation: H.H., K.A., S.W., M.F. and D.K.; Data curation: H.H., K.A. and M.F.; Visualization: H.H. and K.A.; Writing (original draft): K.A.; Writing (review & editing): S.Y., M.F. and D.K.; Supervision: M.F. and D.K.; Project administration: M.F. and D.K.; Funding acquisition: S.Y., T.C., S.H., M.F. and D.K.

All authors contributed to discussions and manuscript preparation.

## Materials and Methods

### Materials

#### *C. elegans* strains

N2 Bristol

CX12725 *inx-20(ok681)*

CB1370 *daf-2(e1370)*

CB502 *sma-2(e502)*

CF1038 *daf-16(mu86)*

YB5013 *inx-20(ok681); daf-2(e1370)*

YB5014 *inx-20(ok681); sma-2(e502)*

YB5661 *inx-20(ok681); daf-16(mu86)*

YB5685 *inx-20(td536)* [*inx-20*::2×G4S::mStayGold::2×G4S::AID*]

BA17 *fem-1(hc17)*

YB4909 *inx-20(ok681); fem-1(hc17)*

YB5274 *inx-20; tdEx2841*[pMF1084.1(5 ng/µl) + pMF1096.2(100 ng/µl)] YB5277 *inx-20; tdEx2844*[pMF1096.2(100 ng/µl)]

### Bacteria strain

*Escherichia coli* OP50

### Plasmids

pMF1084: 5.3kb P*inx-20*::*inx-20* ORF::G4S::mVenus cDNA::*unc-54* 3’UTR

pMF1096: P*myo-3*::mCherry::*unc-54* 3’UTR

“G4S_mStayGold_G4S_AID”

pRF4 (Mello et al., 1991)

### Primers

HH145 To detect *inx-20(ok681)* (Fwd)

GCTCAAGCACTTCCCCAACATTCCG

HH165 To detect *inx-20(ok681)* (Rev)

GCAGATCTATCATAGTCGTCACCG

MF4207: To detect *daf-16(mu86)* deletion (Fwd)

CACCACGACGCAACACACTAATAGTG

MF4208: To detect *daf-16(mu86)* deletion (Rev)

CGTGGTATGATGGTGGTGGAGC

MF3771: To detect *daf-16* inside *mu86* (WT) (Fwd)

aagagcacagacagagtaggggcg

MF3772: To detect *daf-16* inside *mu86* (WT) (Rev) GCGTGGTATGTCTAGACAGTGTGTCCG

KA36 To amplify *inx-20*::G4S::mSG::G4S::AID* repair template (Fwd)

GCGACGGAAAACATCAGTTTTGGTTCCTCTCATGAGTCGAGAAGATTTACATTTGGATCAATCTCCAACTACACCAGCTCCTCAATTCCTTCGtCCaCCtAGCAGtCGtATGGCTTCAGCTGCGAATGTAGGTGGCGGAGGTAGCGGAGGCGGTG

KA37 To amplify *inx-20*::G4S::mSG::G4S::AID* repair template (Rev)

CATTGTTTCCTTTTTCTATGTTGGTTTTATTATTATTTTAATAATTATTATTTCATGTCATGCAAATATTTGAAATTGTAACAACAATTAATTCGGTATCAGGAAAACAACAAAATATTGAAATTATTACTTCACGAACGCCGCCGCCTCCGGGC

KA38 To sequence *inx-20*::G4S::mSG::G4S::AID* insertion (Fwd)

gtgctcatctagttaatgcaaatttgttctgc

KA39 To sequence *inx-20*::G4S::mSG::G4S::AID* insertion (Rev)

CCTTCCTGCTCAAACCATCTAAATACTTCC

## Methods

### Maintenance of *C. elegans*

Worms were cultured as described previously (Brenner, 1974). Unless otherwise noted, worms were maintained at 20 °C on Nematode Growth Medium (NGM) plates seeded with *Escherichia coli* OP50. Strains used in this study are listed in the Materials section.

### Forward genetic screening for mutants which exhibit extended reproductive span

Worms were mutagenized with ethylmethane sulfonate (EMS) as described previously (Brenner, 1974). Worms were maintained on 100 mm 4×peptone plates (Chiyoda *et al*., 2021). L4-stage P_0_ worms were treated with 50 mM EMS, and 7,200 F_1_ animals were obtained. Approximately 12 F_2_ progeny from each F_1_ lineage were screened to recover homozygous mutations. During passaging of the F_2_ generation, worms were collected from plates using M9 buffer (3 g KH_2_PO_4_, 15 g Na_2_HPO_4_·12H_2_O, 5 g NaCl, and 1 ml of 1 M MgSO_4_ dissolved in 1 L H_2_O and autoclaved). Worms were allowed to settle naturally in M9, and the supernatant containing F_3_ larvae and embryos was removed. This procedure was repeated approximately three times, after which the settled F_2_ adults were transferred to fresh plates. F_2_ animals were passaged approximately twice per day. In some cases, worms were kept at 12-15 °C overnight to delay hatching of F_3_ progeny. Strains in which F_3_ progeny were present on plates when F_2_ animals reached adult day 10 or adult day 11 were recovered as candidates with extended reproductive span.

### Next-Generation Sequencing (NGS)

NGS analysis was performed with support from the Platform for Advanced Genome Science. Sequencing reads were mapped to the WBcel235 reference genome. Variants were identified using the Resequencing Analysis tool in CLC Genomics Workbench ver. 23 (QIAGEN). Mutations that differed completely (100%) between the mutant and N2 samples and that either caused amino acid substitutions or were located at splice sites were extracted.

### CRISPR-Cas9 genome editing of *inx-20*

Homology-directed genome editing with CRISPR-Cas9 (Paix *et al*., 2015) was used to fuse mStayGold (Ivorra-Molla *et al*., 2024) to the C-terminal of *inx-20* prior to the stop codon/3′ UTR of *inx-20*. Briefly, the sequence of mStayGold cDNA was optimized using *C. elegans* codon adapter (Redemann *et al*., 2011). 2×G4S::mStayGold::2×G4S::AID* tag was amplified from G4S::mStayGold::G4S::AID* plasmid with KA36, KA37 primer pair resulting in a PCR repair template that contained 130 bp (Fwd) or 125 bp (Rev) of homology to either side of the Cas9 cut site. The C-terminal of *inx-20* was targeted with the crRNA (TCGACCTCCAAGCAGCCGAA), and edits were selected using the pRF4 co-injection (Mello *et al*., 1991). A mix as described previously was injected into the germline of adult *C. elegans* (Dokshin *et al*., 2018). F_1_ progenies were screened for the Rol phenotype and the CRISPR allele was confirmed using PCR amplification with KA38, KA39 primer pair, sequenced, and outcrossed against N2 twice.

### Reproductive span measurement (self-fertilization and mated with male)

30 to 50 L4-stage hermaphrodites per strain were singly placed on 35 mm NGM plates, and this day was designated as adult day 0. P_0_ animals were transferred daily to fresh 35 mm NGM plates, leaving only the embryos laid on that day on the old plates. The presence or absence of F_1_ progeny was assessed two days after removal of the P_0_ animals. When no F_1_ progeny were observed for two consecutive days, egg laying was considered to have ceased on the first of those two days. Animals were censored on the day they were found to have died due to climbing the plate walls, exhibited a bagging phenotype, or for other non–age-related causes. For mating, 120-160 worms in total were placed on a 60 mm plate at a one to three ratio of hermaphroditic L4-stage larvae to N2 adult day one or day two males, and allowed to mate for 24 hours. Progeny in which no males were observed upon hatching were considered to have undergone insufficient mating and were treated as censored.

### Late reproduction measurement

Thirty to fifty L4-stage hermaphrodites were placed together on 35 mm NGM plates, and this day was designated as adult day 0. Until adult day four, animals were maintained as groups and transferred to fresh 35 mm NGM plates either daily or every other day. On adult day five, each animal was singly transferred to a 35 mm NGM plate, and the presence or absence of F_1_ progeny on each plate was assessed on adult day seven.

### Lifespan measurement

Ten L4-stage larvae were placed on each 35 mm plate and transferred daily or every other day. Survival was scored based on the presence or absence of a response to gentle prodding with a pick. Worms that died due to matricide or bursting, or that crawled up the wall of the plate and died, were treated as censored on the day of death.

### Fluorescent microscopy

Worms were mounted on 4% agarose pads prepared with M9 buffer and immobilized with 100 µM sodium azide. Images were captured using Zeiss Axio Imager M2 with a Plan-Apochromat 63×/1.4 oil DIC objective. An Axiocam 506 mono camera was used for image acquisition. mStayGold fluorescence was imaged using 488 nm excitation and intestinal autofluorescence was imaged using 405 nm excitation. Images were acquired using Zeiss Zen software. Brightness and contrast were adjusted using Fiji.

### Dauer induction

Approximately 30-100 fertilized eggs laid on 60 mm plates at 20 °C were transferred to 25 °C. Three days later, the numbers of dauer larvae and adults were counted. Individuals exhibiting dauer alae were scored as dauer larvae, whereas individuals containing fertilized eggs in the uterus were scored as adults.

### Body size measurement

Day one adult worms were mounted on 4% agar pads prepared with M9 buffer and imaged using an S9D stereo zoom microscope equipped with an MC190 HD camera and a KL300 LED light source (LEICA). Body length was measured from the acquired images using WormSizer (Moore, Jordan and Baugh, 2013).

