## Supplementary figures and images for "Gap junctions in the alimentary tract regulate reproductive span in *C. elegans*"

### Supplementary Figures

#### Figure S1

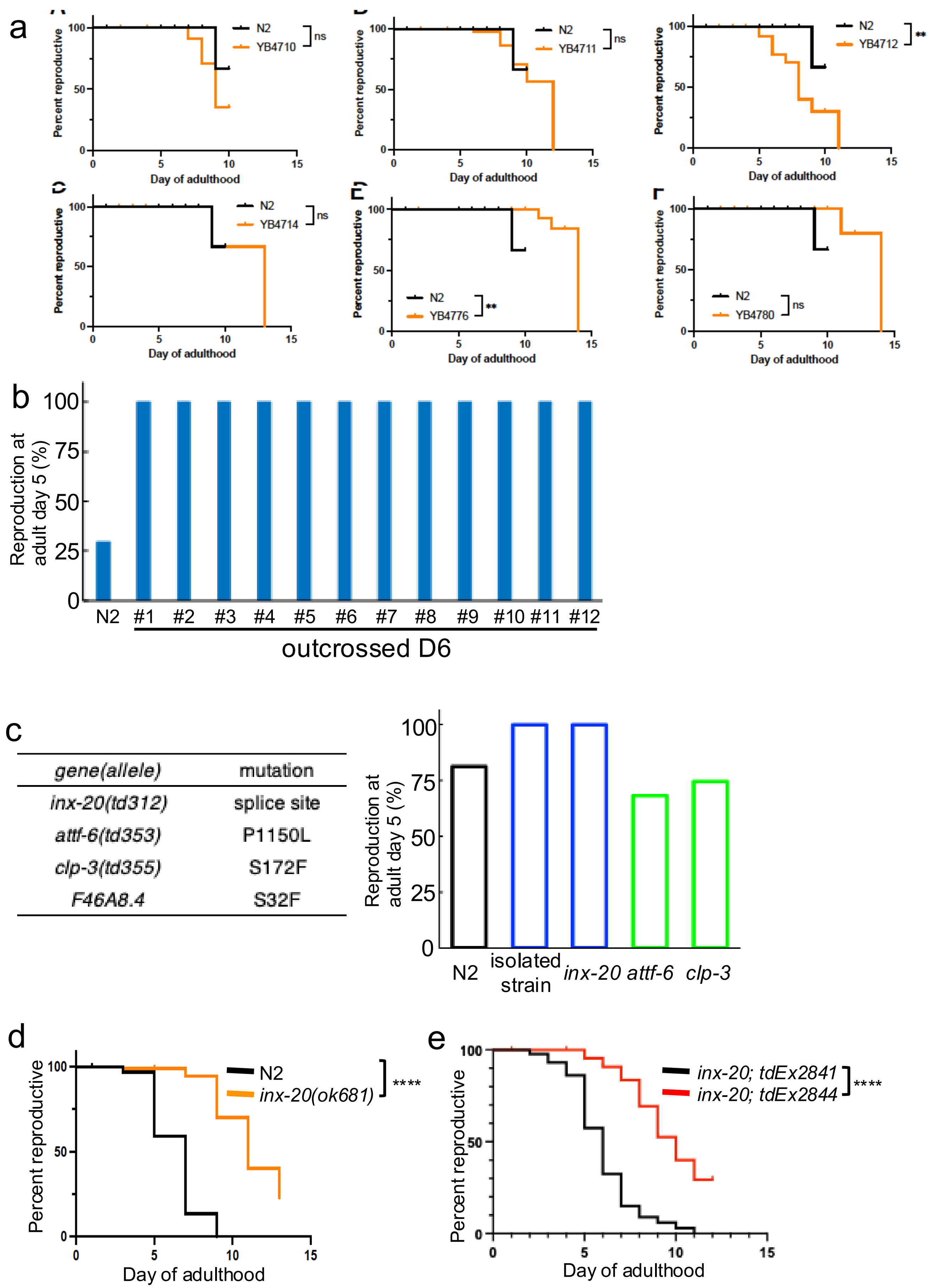


| *gene(allele)* | mutation |
|---|---|
| *inx-20(td312)* | splice site |
| *attf-6(td353)* | P1150L |
| *clp-3(td355)* | S172F |
| *F46A8.4* | S32F |

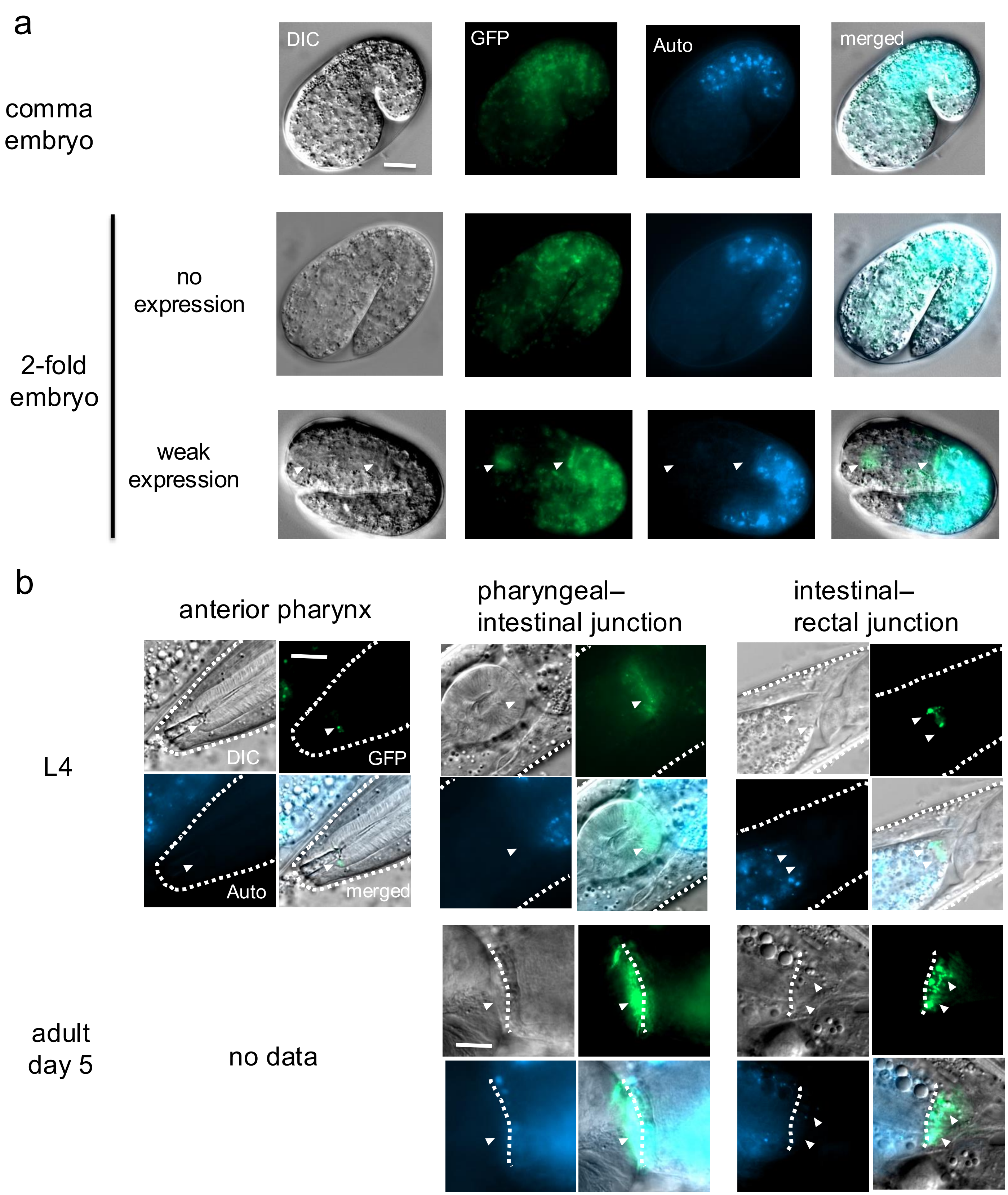
